# Visualizing metabolic network dynamics through time-series metabolomics data

**DOI:** 10.1101/426106

**Authors:** Lea F. Buchweitz, James T. Yurkovich, Christoph M. Blessing, Veronika Kohler, Fabian Schwarzkopf, Zachary A. King, Laurence Yang, Freyr Jóhannsson, Ólafur E. Sigurjónsson, Óttar Rolfsson, Julian Heinrich, Andreas Dräger

## Abstract

New technologies have given rise to an abundance of -omics data, particularly metabolomics data. The scale of these data introduces new challenges for the interpretation and extraction of knowledge, requiring the development of new computational visualization methodologies. Here, we present a new method for the visualization of time-course metabolomics data within the context of metabolic network maps. We demonstrate the utility of this method by examining previously published data for two cellular systems—the human platelet and erythrocyte under cold storage for use in transfusion medicine.

The results comprise two animated videos that allow for new insights into the metabolic state of both cell types. In the case study of the platelet metabolome during storage, the new visualization technique elucidates a nicotinamide accumulation which mirrors that of hypoxanthine and might, therefore, reflect similar pathway usage. This visual analysis provides a possible explanation for why the salvage reactions in purine metabolism exhibit lower activity during the first few days of the storage period. The second case study displays drastic changes in specific erythrocyte metabolite pools at different times during storage at different temperatures.

In conclusion, this new visualization technique introduced in this article constitutes a well-suitable approach for large-scale network exploration and advances hypothesis generation. This method can be applied to any system with data and a metabolic map to promote visualization and understand physiology at the network level. More broadly, we hope that our approach will provide the blueprints for new visualizations of other longitudinal -omics data types.

**AUTHOR SUMMARY:** Profiling the dynamic state of a metabolic network through the use of time-course metabolomics technologies allows insights into cellular biochemistry. Interpreting these data together at the systems level provides challenges that can be addressed through the development of new visualization approaches. Here, we present a new method for the visualization of time-course metabolomics data that integrates data into an existing metabolic network map. In brief, the metabolomics data are visualized directly on a network map with dynamic elements (nodes that either change size, fill level, or color corresponding with the concentration) while the user controls the time series (i.e., which time point is being displayed) through a graphical interface. We provide short videos that illustrate the utility of this method through its application to existing data sets for the human platelet and erythrocyte. The results presented here give blueprints for the development of visualization methods for other time-course -omics data types that attempt to understand systems-level physiology.

## INTRODUCTION

Over the last few decades, new technological developments have enabled the generation of vast amounts of “-omics” data [29]. These various -omic data types have helped bring new insights to a vast array of biological questions [24, 21, 41]. As more and more data are generated, however, researchers are faced with the enormous challenge of integrating, interpreting, and visualizing these data. The community has recognized these needs, focusing efforts on data visualization as a way to maximize the utility of biological data [5]. Data visualization is particularly crucial for a systems-level perspective of metabolic networks and pathways. Several excellent software tools were made available for drawing and exploring biological network graphs [14, 33, 9,10,13]. These tools provide impressive descriptions of the network and support for diverse analyses, including the mapping of omics data to networks.

Metabolomics data provide snapshots of cellular biochemistry, presenting essential insights into a cell’s metabolic state [27, 19]. Visualization tools often allow users to overlay pathway maps with static data sets [14]. Recently, time-course metabolomics data sets that detail cellular changes over time are becoming more prevalent [26, 43, 3, 25], leading to the need for dynamic visualizations that can capture the aspect of time [31]—an essential aspect of understanding complex processes such as changes in metabolic activity, concentration, or availability. Many visualization tools [39, 36, 30, 17], however, do not yet provide support for the representation of dynamic content. Those visualization tools whose features do include time series visualization [5, 28,15, 30,11, 31] only provide static depictions of the data. Some progress has been made to provide a stepwise temporal representation of metabolomics data [1], but a robust and smooth dynamic solution for mapping time series data to networks has yet to be presented.

One reason for the current lack of convincing visual analysis methods for dynamically changing data sets is that time-dependent data add additional layers of complexity to the already difficult problem of visual network exploration. First of all, genome-scale metabolic networks can have enormous sizes: Some published metabolic network maps comprise several thousand biochemical reactions [4, 22], of which human beholders can simultaneously only grasp a very small fraction [12]. Another difficulty in the interpretation of metabolic networks is their small-world property [37]. It means that the connectivity of their nodes (the metabolites) follows a power-law distribution, i.e., a few nodes are highly connected hub-nodes, whereas the majority has only very few connections. Examples for common hub-nodes include currency metabolites such as ATP, NAD(P)H or cofactors. Overall, nodes are connected through very few consecutive edges (the reactions). Furthermore, metabolic networks are hypergraphs, in which more than one metabolite can act as reactant or metabolites, i.e., one edge may connect multiple nodes at once. To circumvent this problem, metabolic networks are often displayed in form of bipartite graphs, introducing a specific reaction node type to which metabolites nodes are connected while prohibiting edges between nodes of the same type [23, 14, 18]. Common graph drawing algorithms, however, usually do not take this hypergraph or bipartite property into account. Very large networks with multiple thousand nodes may, therefore, result in hairball-like structures, centered around hub-nodes. Duplicating selected hub-nodes has been suggested as a strategy for reducing the problem of hairball formation [18] and is commonly used in manually-drawn networks because it leads to significantly better indication of conceptual sub-maps and flows of matter [22]. However, this technique introduces another problem when mapping data onto the network. The same node is displayed in varying location, therefore also is the quantitative value associated to that node. Particularly in dynamically changing graphs, drastic alterations in duplicated nodes may distract beholders from less obvious activities in the network. Moreover, metabolic networks traditionally comprise characteristic structures, in particular cycles such as the tricarboxylic acid (TCA) cycle or the urea cycle. Not only can a node duplication in the wrong position destroy the circular shape of this graph pattern, but algorithmically drawn networks without such prior knowledge may not consider structures that researchers usually expect because of their familiarity with common textbook representations. For these reasons, automatically drawing a metabolic network from scratch and mapping data to it without human interaction is unlikely produce convincing results, nor can it directly identify hidden activities.

With a steadily increasing number of carefully prepared metabolic network layouts being published, we here assume a map to be available for the system of interest. If this is not yet the case, a map can be easily drawn using software such as Escher [14]. This paper focuses on the problem of displaying dynamically changing quantitative data of network components. The aim is to answer the question: How to create expressive visual displays of dynamic metabolic networks? Needed are strategies to visually present the data in a way that beholders can best perceive and estimate quantities of network individual components and that at the same time enable them to conceptually narrow down parts of interest even within large networks.

In the next sections, we present an intuitive and comprehensive method for the visualization and contextualization of longitudinal metabolomics data in metabolic networks. We developed three different graphical representations of metabolic concentration that allow for different interpretations of metabolomics data through a smooth animation. We present two case studies using this method that examine two different cellular systems—the human platelet and the human red blood cell (RBC)—to show how visualizing existing data can provide new insights into cellular metabolism. The result are two animated videos that give detailed information about the systems under study and highlight new insights that were not previously apparent. We hope that this intuitive method will aid researchers in interpreting and visualizing complex data sets.

## RESULTS

In this study, we present a new approach for the visualization of time-course metabolomics data in the context of large-scale metabolic network maps. The idea is that time series can be adequately observed in the form of an animated sequence of a dynamically changing network map when using an appropriate representation of metabolic quantities. To this end, our technique exploits the repeatedly observed ability of human beholders to estimate quantities most precisely when these are mapped to a lengths scale [6]. Since metabolic maps commonly represent nodes with circles [16,14], we suggest using the fill level of each node as a visual element to represent its amount at each time point. We experimented with visualization of data in several different ways, based on node size, color, a combination of size and color, or fill level (Supplementary Figure S1). Each of these visual representations provides some advantages over the others, but the notion of the fill level of a node can be the most intuitive [6] as it allows for the user to understand and gauge its minimum or maximum value quickly (see Discussion).

Using this technique, we created such an animation for given longitudinal metabolomics data and a metabolic network map that corresponds to the observed cell type (Figure 1). To provide a smooth animation, additional time points are interpolated in the provided time series. Further details regarding the development and use of the implementation of the method can be found in the Supplementary Information.

**Fig. 1.**
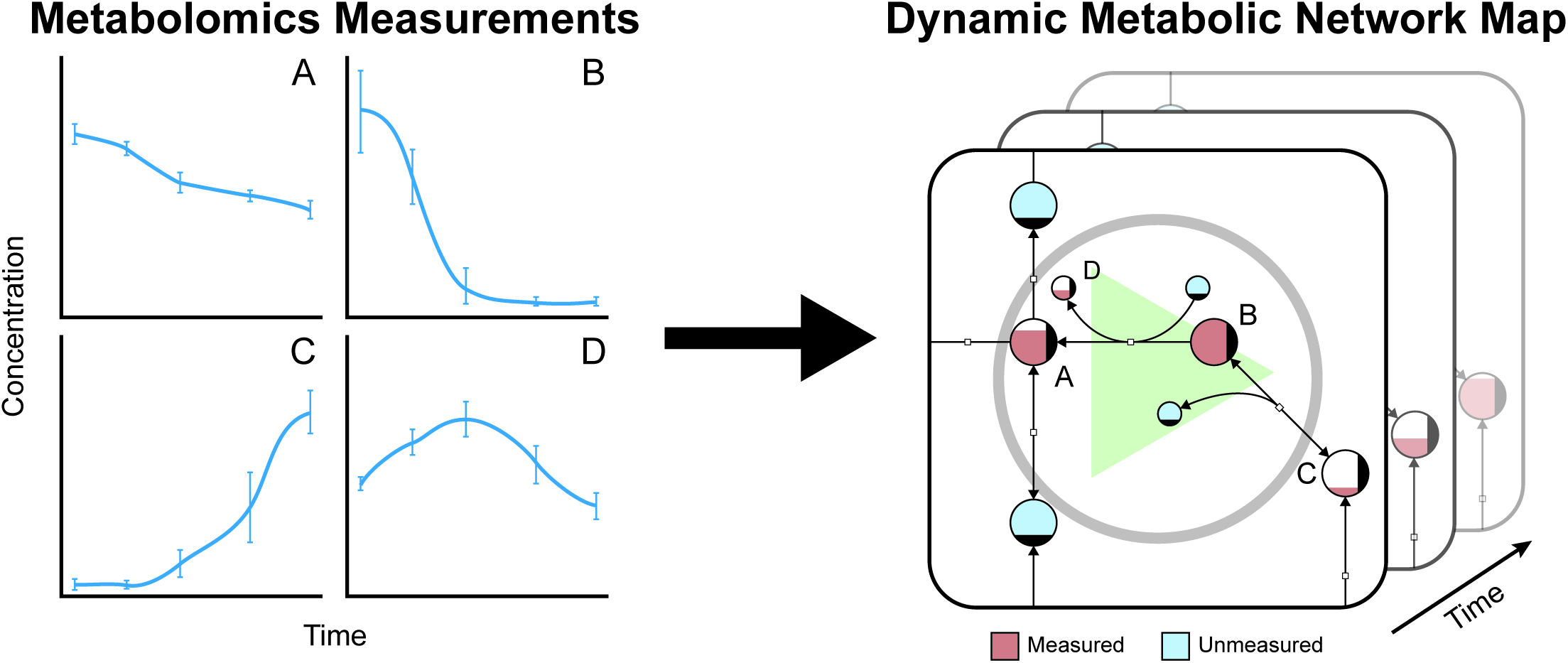
Dynamic visualization of metabolomics data. We take metabolomics data as input and generates a dynamic animation of the data over time which enables the visualization of pool sizes for individually measured metabolites. Several different options are discussed in this article for the visualization of the data based on node size, color, and fill level. The method has been implemented in SBMLsimulator including an export function to save the resulting output in a video file. For creation of animation videos highlighted in Box 1 and Box 2 post-processing steps are needed as descibed in the Supplementary Information.

To demonstrate the utility of this method, we applied these visualization methods to two different cellular systems—human platelets and RBCs—for which longitudinal quantitative data sets were available in the literature [26, 43]. Transfusion medicine plays a vital role in modern healthcare, making the storage of different blood components important physiological processes to understand. In particular, platelets and RBCs represent relatively simple human cell types that can be intensely studied in the well-defined, static environment provided by blood storage (packed in plastic bags and stored at 22 °C and 4 °C for platelets and RBCs, respectively). While the cells are stored in these conditions, biochemical and morphological changes occur (the “storage lesion”) that are well-studied through the use of metabolomics data [19, 40]. Metabolic models were previously available for both the platelet [34] and RBC [2], enabling the creation of network maps for both reconstructions. Thus, these data could be visualized in the context of the entire metabolic network.

### Case study: human platelets under storage conditions

Our first case study examined the storage of platelets. We manually created a metabolic map for the complete metabolic network of the platelet using Escher [14]. We then overlaid metabolomics data which characterized the baseline storage conditions with eight time points over ten days of storage [26] to produce a network-level visualization of the data (Figure 2). Using this network-level visualization, we examined the dynamics of the platelet metabolome.

**Fig. 2.**
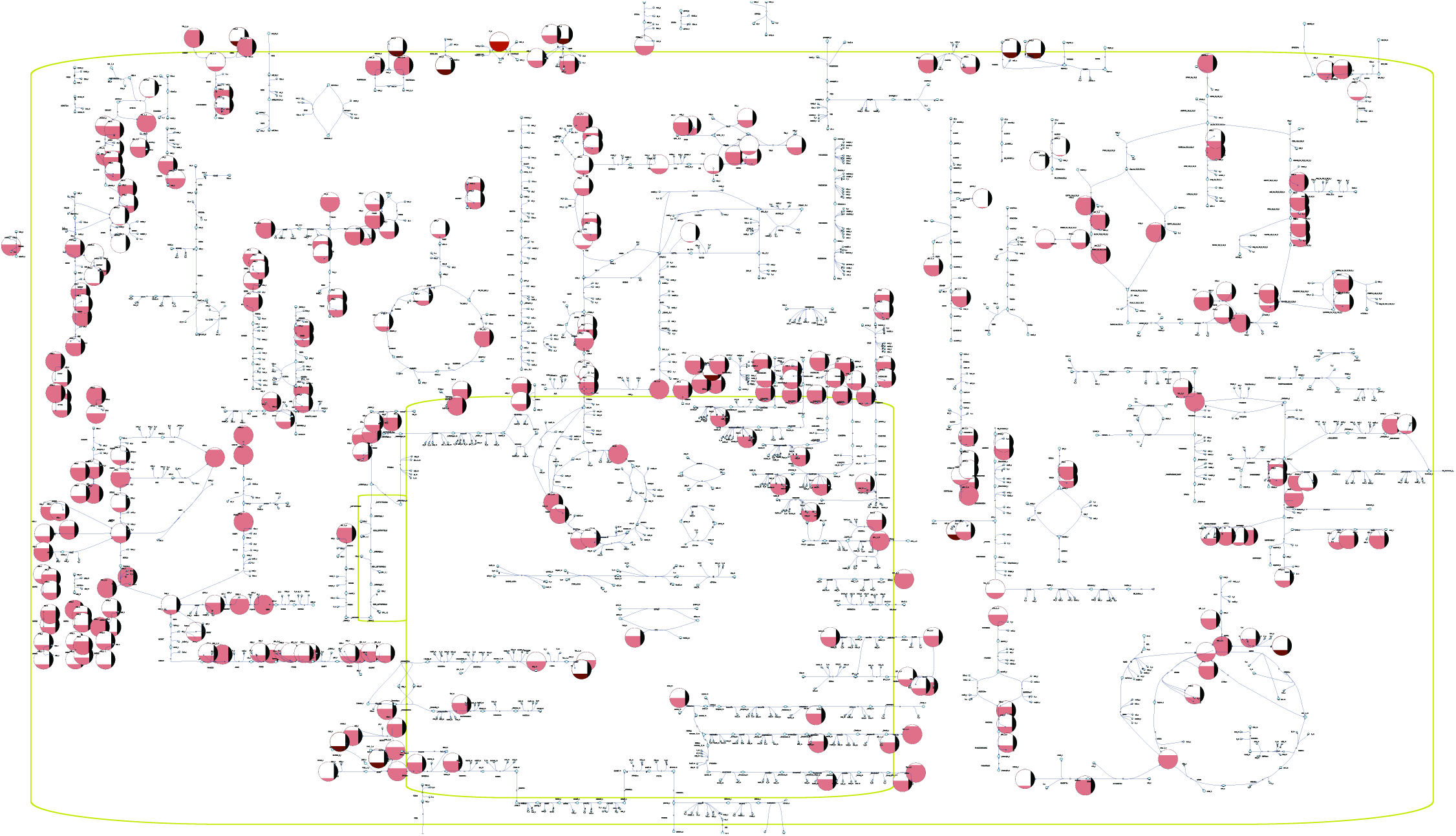
Network map in SBGN style [16] for the human platelet with metabolomics data [26] overlaid. This figure represents a visualization in which the fill level of a node represents the relative size of the corresponding metabolite pool.

During the first part of storage, stress due to the non-physiological conditions of storage (i.e., packed in a plastic bag at 22 °C) slows metabolic activity through glycolysis, the pentose phosphate pathway, and purine salvage pathways [26]. Several metabolites are secreted by the cells and accumulate in the storage media, such as hypoxanthine. The metabolite 5-Phospho-alpha-D-ribose 1-diphosphate (PRPP) is produced from the pentose phosphate pathway and is a cofactor in the salvage reactions that break down hypoxanthine. Because flux through the pentose phosphate pathway is lower, the cells have less capacity to recycle hypoxanthine using the salvage pathways.

##### Box 1: Visualization of Biochemical Processes - Storage of Platelets © 8 min: 26 s

This video introduces a new method for visualizing metabolic processes in the context of a full biochemical network. Representing the metabolic network as a graph where metabolites are nodes and reactions are edges can help elucidate complex relationships within the network. While viewing a network in this manner is not new, overlaying -omics data onto the map allows for an accurate integration of disparate data types. By visually interpreting the information in this dynamic, graphical format, we can more easily distinguish important characteristics of the network. This video utilizes the metabolomics data from the study “Comprehensive metabolomic study of platelets reveals the expression of discrete metabolic phenotypes during storage” [26].

By viewing all of the data simultaneously at the network level, we were able to discover that the concentration profile of nicotinamide mirrors that of hypoxanthine. This observation suggests a similar rationale for the accumulation of nicotinamide, providing a hypothesis as to why the salvage pathway within purine metabolism has lower activity during the first few days of storage. These findings are demonstrated in the video highlighted in (Box 1), helping show how network-level visualization allows for improved extraction of biological insight from large, complex data sets.

### Case study: human red blood cells under storage conditions

Our second case study examined the storage of RBCs. A metabolic map was already available for the RBC [42] and captures the complete metabolic network [2]. Here, we sought to examine a data set that provided the opportunity to visualize different experimental conditions for the same network. Recently, a study was published [43] that used quantitative longitudinal metabolomics data to examine the state of the RBC metabolome under four different storage temperatures: 4 °C (storage temperature), 13 °C, 22 °C, and 37 °C (body temperature). For this system, we opted to visualize the dynamics of the metabolite concentrations as nodes with variable size where smaller nodes represent smaller pool sizes, and larger nodes represent larger pool sizes (Figure 3).

**Fig. 3.**
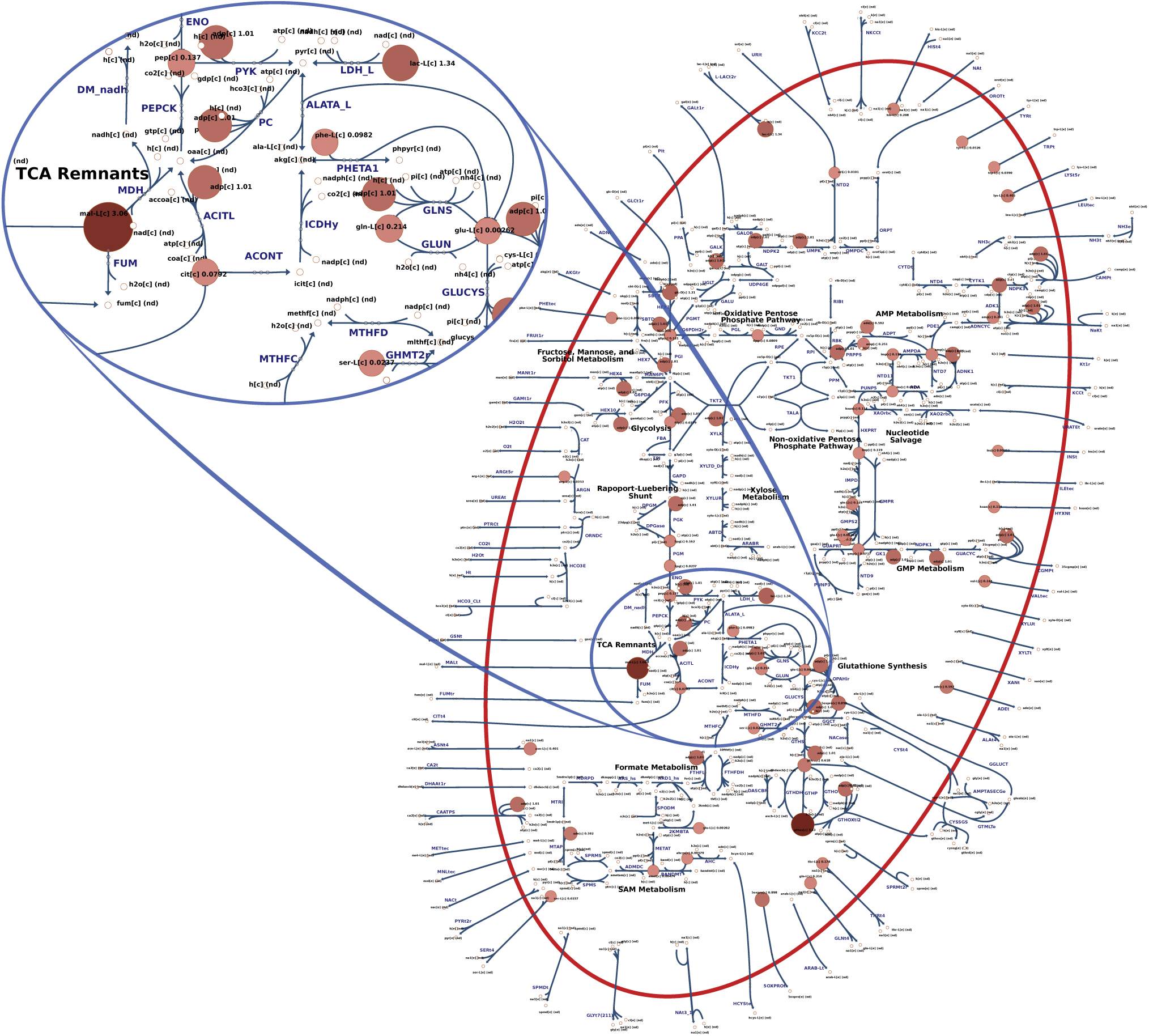
Overview of the RBC metabolic network under storage conditions at 4 °C. The size and color of the nodes reflect their absolute abundance. The oval area on the top magnifies a region in the center of the map that appears in the style of Escher [14] in contrast to the SBGN style shown in Figure 2.

To highlight the differences between the experimental conditions, we examined two of the conditions side-by-side (see the video highlighted in Box 2). This visualization helps supplement the type of statistical and modeling analyses performed previously and helps contextualize the effects of the temperature change across different parts of the network. In particular, it is obvious from a network-level view of the system that certain parts of the network are more active at different points in the time-course. A side-by-side comparison helped emphasize that the availability of reduced glutathione is different with increased temperature, an important physiological feature due to the role of glutathione in neutralizing reactive oxygen species [38] that accumulate during storage and contribute to the storage lesion [7]. Finally, it can be seen that hypoxanthine—a known toxic metabolite whose concentration has been shown to inversely correlate with the post-transfusion recovery rates of transfusion patients [20]—accumulates faster at higher temperatures. Like in the other case study presented above, the new insights into complex processes (which are not yet fully understood) provide evidence that this method can be beneficial for the simplification and understanding of large, complex data analyses.

##### Box 2: Visualization of Biochemical Processes - Temperature Dependence of Red Blood Cells © 1 min: 33 s

This video visually compares the biochemical effects of increasing the storage temperature from (4 °C to 13 °C) of stored RBCs on metabolic processes. The relative node size shows changes in metabolite concentrations for each measured metabolite. Zooming in on various parts of the network helps visualize how specific metabolite pools undergo more drastic changes at different points during storage. This video utilizes the metabolomics data from the study “Quantitative time-course metabolomics in human red blood cells reveal the temperature dependence of human metabolic networks” [43].

## DISCUSSION

With the development of new experimental technologies and the subsequent generation of -omics data sets, life scientists are faced with the challenge of extracting actionable knowledge. New visualization methods are a critical way that the community can make strides toward making the most of complex data. Here, we present a new method for the visualization of longitudinal metabolomics data in the context of the metabolic network. We provide two case studies that examine (1) a baseline characterization of a physiological process and (2) a set of experimental perturbations that allowed for a side-by-side comparison of different experimental conditions. The introduction of this new visualization method has two significant implications.

First, this method provides a dynamic visualization of cellular processes. Tools such as Cytoscape [32] provide visual analysis of networks and supports plugins like TiCoNE [39] and CyDataSeries [35] for the visualization of time-course data. However, tools such as these or VANTED [30] only offer static representations of dynamic data. To our knowledge, only KEGGanim [1] offers a dynamic visualization of time-course data. The method presented here builds on KEGGanim by offering a smooth interpolation between time points and offers the further advantage of customization concerning the display of both data and the network itself. The method presented outlines an original development for visualizing complex biological data in an intuitive and useful manner.

Second, a network-level representation of large metabolomics data sets presents a more holistic view of the data than does statistical analysis alone. While visual inspection of data is indeed not a replacement for more detailed statistical or modeling analyses, this method provides an important supplement to existing data analysis pipelines. We demonstrate its utility in such an analysis pipeline by highlighting findings from existing data sets [43, 26]. Visualizing the metabolomics data in the context of the full metabolic network allowed for new insights into existing data sets. A potential explanation why the salvage pathway lowers its activity during the first few days of platelet storage could be deduced for the network of the human platelet. In the RBC network, it could easily be seen that concentrations in certain parts of the network (e.g., nucleotide metabolism) accumulated or depleted together. These findings illustrate the promising potential of intuitively visualized time-course data and—combined with in-depth computational data analysis—can help elucidate physiological processes.

The simplification of experimental data interpretation became extremely relevant in the age of high-throughput technologies. The visualization concept presented here offers an intuitive, systems-level interpretation of metabolomics data. Combined with other data analytics, this method helps provide a holistic view of a data set, moving us closer to being able to realize the full potential of a given data set. More broadly, we hope that the method presented here will provide the starting point for further visualization improvements not only for metabolomics data but for the visualization and contextualization of other data types. Future work may include combining a dynamic representation with static concentration graphs that will continue to improve the capabilities of such software to fully meet the needs of life science researchers.

## METHODS

The method described in this paper utilizes existing software libraries to visually represent metabolomics data in the context of a metabolic network map. SBMLsimulator [8] is used to interpolate the data, providing a smooth time-course for simulation. Escher [14] is used for the design of metabolic network maps. In brief, metabolomics data are provided in a `*.csv` file format with identifiers matching those of the map. The data are interpolated over time with input from the user. Other features are selected, such as the speed of animation and how metabolite concentrations are represented (e.g., fill level). The result is a smooth animation that allows features such as zooming and panning across different areas of the map. Full details for the implementation and use of the software are provided in the Supplemental Material.

## ACKNOWLEDGMENTS

The authors would like to thank Prof. Dr. Robert Feil, Katrin Keppler for her music contribution, Jan D. Rudolph, and Jakob Matthes for implementing a basic Java™ interpreter of SBML layout.

This work was funded by the NIH (R01-GM070923 and U01-GM102098), the Landspitali University Hospital Research Fund, the University of Iceland Research Fund, and the Novo Nordisk Foundation through the Center for Biosustainability at the Technical University of Denmark (NNF10CC1016517).

This work was made possible by the friendly support of yWorks GmbH (https://www.yworks.com) who provided their diagram visualization library yFiles for Java (https://www.yworks.com/products/yfiles-for-java) and assistance during the implementation phase.

## REFERENCES

[1] Adler, P., J. Reimand, J. Jänes, R. Kolde, H. Peterson, et al,, 2008 KEGGanim: pathway animations for high-throughput data. Bioinformatics 24: 588–590.

[2] Bordbar, A., N. Jamshidi, and B. O. Palsson, 2011 iAB-RBC-283: A proteomically derived knowledge-base of erythrocyte metabolism that can be used to simulate its physiological and patho-physiological states. BMC Systems Biology 5: 110.

[3] Bordbar, A., J. T. Yurkovich, G. Paglia, O. Rolfsson, O. E. Sigurjónsson, et al., 2017 Elucidating dynamic metabolic physiology through network integration of quantitative timecourse metabolomics. Scientific Reports 7: 1–12.

[4] Brunk, E., S. Sahoo, D. C. Zielinski, A. Altunkaya, A. Dräger, et al,, 2018 Re-con3D enables a three-dimensional view of gene variation in human metabolism. Nature Biotechnology pp. 1–37.

[5] Callaway, E., 2016 The visualizations transforming biology. Nature 535: 187–188.

[6] Cleveland, W. S. and R. McGrill, 1984 Graphical Perception: Theory, Experimentation and Application to the Development of Graphical Methods. Journal of the American Statistical Association 79: 531–554.

[7] D’Alessandro, A., G. M. D’Amici, S. Vaglio, and L. Zolla, 2012 Time-course investigation of SAGM-stored leukocyte-filtered red bood cell concentrates: from metabolism to proteomics. Haematologica 97: 107–115.

[8] Dörr, A., R. Keller, A. Zell, and A. Dräger, 2014 SBMLSimulator: A Java Tool for Model Simulation and Parameter Estimation in Systems Biology. Computation 2: 246–257.

[9] Droste, P., K. Nöh, and W. Wiechert, 2013 Omix – a visualization tool for metabolic networks with highest usability and customizability in focus. Chem. Ing. Tech. 85: 849–862.

[10] Funahashi, A., Y. Matsuoka, A. Jouraku, M. Morohashi, N. Kikuchi, et al,, 2008 CellDesigner 3.5: A versatile modeling tool for biochemical networks. Proc. IEEE 96: 1254–1265.

[11] Gehlenborg, N., S. I. O’Donoghue, N. S. Baliga, A. Goesmann, H. Hibbs, M. A. Kitano, et al,, 2010 Visualization of omics data for systems biology. Nature Methods 7: S56–S68.

[12] Halford, G. S., R. Baker, J. E. McCredden, and J. D. Bain, 2005 How many variables can humans process? Psychological Science 16: 70–76.

[13] Kelder, T., M. P. van Iersel, K. Hanspers, M. Kutmon, B. R. Conklin, et al., 2012 WikiPathways: building research communities on biological pathways. Nucleic Acids Res. 40: D1301–7.

[14] King, Z. A., A. Dräger, A. Ebrahim, N. Sonnenschein, N. E. Lewis, et al., 2015 Escher: A web application for building, sharing, and embedding Data-Rich visualizations of biological pathways. PLoS Comput. Biol. 11: e1004321.

[15] König, M. and H.-G. Dräger, Andreas Holzhütter, 2012 CySBML: a Cytoscape plugin for SBML. Bioinformatics 28: 2402–2403.

[16] Le Novère, N., M. Hucka, H. Mi, S. Moodie, F. Schreiber, et al., 2009 The Systems Biology Graphical Notation. Nature Biotechnology 27: 735–741.

[17] Ma, D. K. G., C. Stolte, S. Kaur, M. Bain, and S. I. O’Donoghue, 2013 Visual analytics of phosphorylation time-series data on insulin response. AIP Conference Proceedings 1559: 185–196.

[18] Moodie, S., N. Le Novère, E. Demir, H. Mi, and A. Villeger, 2015 Systems Biology Graphical Notation: Process Description language Level 1 Version 1.3. Journal of integrative bioinformatics 12: 263.

[19] Nemkov, T., K. C. Hansen, L. J. Dumont, and A. D’Alessandro, 2016 Metabolomics in transfusion medicine. Transfusion 56: 980–993.

[20] Nemkov, T., K. Sun, J. A. Reisz, A. Song, T. Yoshida, et al., 2017 Hypoxia modulates the purine salvage pathway and decreases red blood cell and supernatant levels of hypoxanthine during refrigerated storage. Haematologica.

[21] Nielsen, J., 2017 Systems Biology of Metabolism. Annual Review of Biochemistry 86: 245–275.

[22] Noronha, A., A. D. Daníelsdóttir, P. Gawron, F. Jóhannssón, S. Jónsdottir, et al., 2017 Reconmap: an interactive visualization of human metabolism. Bioinformatics 33: 605–607.

[23] Noronha, A., P. Vilaça, and M. Rocha, 2014 An integrated network visualization framework towards metabolic engineering applications. BMC Bioinformatics 15: 1–13.

[24] Österlund, T., M. Cvijovic, and E. Kristiansson, 2017 Integrative Analysis of Omics Data. In Systems Biology, edited by J. Nielsen and S. Hohmann, chapter 1, pp. 1–24, Wiley-VCH Verlag GmbH & Co. KGaA, Weinheim.

[25] Paglia, G., Ó. E. Sigurjonsson, Ó. Rolfsson, M. B. Hansen, S. Brynjólfsson, et al., 2015 Metabolomic analysis of platelets during storage: a comparison between apheresis- and buffy coat-derived platelet concentrates. Transfusion 55: 301–313.

[26] Paglia, G., O. E. Sigurjónsson, O. Rolfsson, S. Valgeirsdottir, M. B. Hansen, et al., 2014 Comprehensive metabolomic study of platelets reveals the expression of discrete metabolic phenotypes during storage. Transfusion 54: 2911–2923.

[27] Patti, G. J., O. Yanes, and G. Siuzdak, 2012 Innovation: Metabolomics: the apogee of the omics trilogy. Nature reviews. Molecular cell biology 13: 263–269.

[28] Pavlopoulos, G. A., A.-L. Wegener, and R. Schneider, 2008 A survey of visualization tools for biological network analysis. BioData Mining 1.

[29] Robinson, J. L. and J. Nielsen, 2016 Integrative analysis of human omics data using biomolecular networks. Mol. BioSyst. 12: 2953–2964.

[30] Rohn, H., A. Junker, A. Hartmann, E. Grafahrend-Belau, H. Treutler, et al., 2012 VANTED v2: a framework for systems biology applications. BMC Systems Biology 6.

[31] Secrier, M. and R. Schneider, 2014 Visualizing time-related data in biology, a review. Briefing in Bioinformatics 15: 771–782.

[32] Shannon, P., A. Markiel, O. Ozier, N. S. Baliga, J. T. Wang, et al., 2003 Cytoscape: A Software Environment for Integrated Models of Biomolecular Interaction Networks. Genome Research 13: 2498–2504.

[33] Smoot, M. E., K. Ono, J. Ruscheinski, P.-L. Wang, and T. Ideker, 2011 Cytoscape 2.8: new features for data integration and network visualization. Bioinformatics 27: 431–432.

[34] Thomas, A., S. Rahmanian, A. Bordbar, B. O. Palsson, and N. Jamshidi, 2014 Network reconstruction of platelet metabolism identifies metabolic signature for aspirin resistance. Scientific Reports 4: 1–10.

[35] Černý, M., 2017 a CyDataSeries - Add time series and the like to your networks. http://apps.cytoscape.org/apps/cydataseries.

[36] Černý, M., 2017b Improve handling of time series and similar in Cytoscape. https://github.com/nrnb/GoogleSummerOfCode/issues/76.

[37] Wagner, A. and D. A. Fell, 2001 The small world inside large metabolic networks. Proceedings of the Royal Society B: Biological Sciences 268: 1803–1810.

[38] Whillier, S., J. E. Raftos, R. L. Sparrow, and P. W. Kuchel, 2011 The effects of long-term storage of human red blood cells on the glutathione synthesis rate and steady-state concentration. Transfusion 51: 1450–1459.

[39] Wiwie, C., A. Rauch, A. Haakonsson, I. Barrio-Hernandez, B. Blagoev, et al., 2017 Elucidation of time-dependent systems biology cell response patterns with time course network enrichment. Manuscript sumbitted for publication.

[40] Yurkovich, J. T., A. Bordbar, Ó. E. Sigurjonsson, and B. O. Palsson, 2018 Systems biology as an emerging paradigm in transfusion medicine. BMC Systems Biology 12.

[41] Yurkovich, J. T. and B. O. Palsson, 2018 Quantitative -omic data empowers bottom-up systems biology. Current Opinion in Biotechnology 51: 130–136.

[42] Yurkovich, J. T., B. J. Yurkovich, A. Dräger, B. O. Palsson, and Z. A. King, 2017a a A padawan programmer’s guide to developing software libraries. Cell Systems 5: 431–437.

[43] Yurkovich, J. T., D. C. Zielinski, L. Yang, G. Paglia, O. Rolfsson, et al., 2017b b Quantitative time-course metabolomics in human red blood cells reveal the temperature dependence of human metabolic networks. Journal of Biological Chemistry 292: jbc.M117.804914.

